# Fronto-striatal projections regulate approach-avoidance conflict

**DOI:** 10.1101/2020.03.05.979708

**Authors:** Adrienne C. Loewke, Adelaide R. Minerva, Alexandra B. Nelson, Anatol C. Kreitzer, Lisa A. Gunaydin

**Affiliations:** Institute for Neurodegenerative Diseases, University of California San Francisco, San Francisco CA; Department of Neurology, University of California San Francisco, San Francisco CA; Kavli Institute for Fundamental Neuroscience, University of California San Francisco, San Francisco CA; Department of Physiology, University of California San Francisco, San Francisco CA; Gladstone Institute of Neurological Disease, University of California San Francisco, San Francisco CA; Department of Psychiatry, University of California San Francisco, San Francisco CA

## Abstract

The dorsomedial prefrontal cortex (dmPFC) has been linked to approach-avoidance behavior and decision-making under conflict, key neural computations thought to be altered in anxiety disorders. However, the heterogeneity of efferent prefrontal projections has obscured identification of the specific top-down neural pathways regulating these anxiety-related behaviors. While the dmPFC-amygdala circuit has long been implicated in controlling reflexive fear responses, recent work suggests that this circuit is less important for avoidance behavior. We hypothesized that dmPFC neurons projecting to the dorsomedial striatum (DMS) represent a subset of prefrontal neurons that robustly encode and drive approach-avoidance behavior. Using fiber photometry recording during the elevated zero maze (EZM) task, we show heightened neural activity in prefrontal and fronto-striatal projection neurons, but not fronto-amydalar projection neurons, during exploration of the anxiogenic open arms of the maze. Additionally, through pathway-specific optogenetics we demonstrate that this fronto-striatal projection preferentially excites postsynaptic D1 receptor-expressing medium spiny neurons in the DMS and bidirectionally controls avoidance behavior. We conclude that this striatal-projecting subpopulation of prefrontal neurons regulates approach-avoidance conflict, supporting a model for prefrontal control of defensive behavior in which the dmPFC-amygdala projection controls reflexive fear behavior and the dmPFC-striatum projection controls anxious avoidance behavior. Our findings identify this fronto-striatal circuit as a valuable therapeutic target for developing interventions to alleviate excessive avoidance behavior in anxiety disorders.

## INTRODUCTION

Avoiding danger is a fundamental behavior required for survival. However, animals can receive conflicting external cues that indicate both potential risk (inducing avoidance) and potential reward (inducing approach). To resolve this approach-avoidance conflict, the animal must decide how to proceed based on these opposing inputs. One theoretical framework for the resolution of this conflict is reinforcement sensitivity theory, which involves three opposing systems: the behavioral activation system (BAS), which responds to potential rewards; the fight/flight system (FFS), which responds to imminent threats; and the behavioral inhibition system (BIS), which responds to conflicting drives toward a goal (via BAS) and away from it (via FFS) (1, 2). According to this theory, activation of the BIS leads to a risk-assessment period or delay in action selection, during which more external information can be received (3, 4). While this response is generally adaptive, it can shift toward a maladaptive overestimation of potential threats in individuals with anxiety disorders (5)—an overactivated BIS leads to excessive risk assessment (e.g., hypervigilance, rumination) and persistent avoidance that can produce severe psychosocial impairment. Compared to our mechanistic understanding of reflexive defensive behaviors such as freezing, little is known about the neural circuit dynamics underlying approach-avoidance conflict, representing a major gap in our understanding of anxiety disorders. Identifying the neural circuits underlying avoidance behaviors is critical for developing more targeted symptom-specific treatments.

While reinforcement sensitivity theory offers a conceptual resolution of how approach-avoidance conflict may be resolved, it lacks a concrete mapping onto specific brain circuits. The BIS is fundamentally a decision-making system, with inputs from the surrounding environment and outputs that delay action selection. One candidate neural structure for this function is the medial prefrontal cortex (mPFC), which has been implicated in decision-making (6, 7), cost-benefit analysis (8, 9), and goal-directed actions (10-13)—all central components of the response to approach-avoidance conflict. Additionally, the mPFC receives contextual and valence information (e.g., from the amygdala and hippocampus) (14-16) and projects to downstream basal ganglia targets involved in movement and action selection (17, 18), making it well-situated to directly control avoidance actions based on environmental cues. In rodents, the mPFC is divided into two subregions thought to play opposing roles in defensive behaviors. Dorsomedial PFC (dmPFC), or prelimbic cortex, is implicated in fear expression (19-21), whereas ventromedial PFC (vmPFC), or infralimbic cortex, is implicated in fear extinction (22-24). Additionally, altered prefrontal activity has been associated with anxiety disorders (25-27), and rodent *in vivo* electrophysiological recordings have shown that single units within the mPFC represent aspects of approach-avoidance tasks (28). However, the mPFC is a highly heterogenous region with many downstream targets, making it difficult to identify which projection-defined mPFC subpopulations are causally involved in approach-avoidance behavior. While activity in the dmPFC-amygdala projection has long been associated with fear expression, optogenetic modulation of this circuit has no effect on approach-avoidance behavior (29), suggesting the involvement of an alternative dmPFC projection.

One such potential dmPFC target is the striatum, which controls movement and action selection through two subpopulations of medium spiny neurons (MSNs): direct-pathway MSNs expressing D1-type dopamine receptors that promote movement, and indirect-pathway MSNs expressing D2-type dopamine receptors that inhibit movement. Ventral and dorsomedial aspects of striatum receive prominent innervation from the dmPFC (30, 31) and form basal ganglia circuits that are involved in cognitive/affective behaviors (32, 33). Experimental human studies have found that deep brain stimulation (DBS) of striatal areas can alleviate anxiety symptoms (34, 35), a finding replicated in rodent work (36). Previous studies investigating the role of the striatum in anxiety disorders have primarily focused on the ventral striatum for its role in affective processing (37-39), whereas the dorsomedial striatum (DMS) has traditionally been implicated in locomotion (40). However, the DMS also plays an important role in regulating reinforcement (41, 42), decision-making (43), and several types of avoidance behavior (44-48); notably, the dmPFC-DMS circuit is involved in decision-making under conflict (49), a key component of the risk-assessment basis of approach-avoidance behavior. In a human approach-avoidance conflict task, conflict trials elicited greater caudate (DMS in rodents) activation than non-conflict trials (47). Recently, DMS D2 MSNs were shown to control approach-avoidance behavior (44).

Despite separate lines of evidence that the dmPFC and the DMS are relevant to anxiety and avoidance behavior, no studies have directly examined the role of dmPFC inputs to the DMS in modulating that behavior. Here, we test the importance of this fronto-striatal circuit in approach-avoidance behavior using a combination of optical circuit-dissection techniques to both record (via fiber photometry) and manipulate (via optogenetics) the neural activity of this projection during the elevated zero maze (EZM) task, which measures innate avoidance of risky anxiogenic environments by quantifying the amount of time animals explore ‘open arms’ (exposed and brightly lit platforms with greater risk of predation) compared to the safer ‘closed arms’ with walls. Additionally, we use slice electrophysiology combined with pathway-specific optogenetics to address how dmPFC inputs influence the activity of downstream striatal neurons. These studies highlight the importance of the dmPFC-DMS fronto-striatal circuit in encoding and controlling anxiety-related behaviors.

## METHODS AND MATERIALS

### Animal Subjects

We used wild-type C57BL/6J (Jackson), Tg(Drd1a-cre)EY217Gsat (Jackson), Tg(Adora2a-cre)KG139Gsat (Jackson), and Drd1a-tdTomato mice (50), all on a C57BL/6J background. Animals were raised in normal light conditions (12:12 light/dark cycle), fed and watered *ad libitum*. All experiments were conducted in accordance with procedures established by the Institutional Animal Care and Use Committee at the University of California, San Francisco.

### Stereotaxic Surgery, Viral Injections, and Fiber Optic Cannula Implantation

Surgeries were performed at 10-14 weeks of age. For fiber photometry, we injected 500 nL of AAV5-CaMKII-GCaMP6f into the dmPFC to record pyramidal neuron activity; to record dmPFC-DMS and dmPFC-BLA projection neurons, we injected 1500 nL of AAV1-Syn-Flex-GCaMP6m into the dmPFC and either 350 nL each of CAV2-Cre and hSyn-mCherry in the DMS or 250 nL each in the BLA. For all fiber photometry experiments, we implanted a 2.5 mm metal fiber optic cannula with 400 µm fiber optic stub (Doric Lenses) in the dmPFC and waited 4-5 weeks for viral expression. For dmPFC cell body and projection optogenetic experiments, we injected either 500 nL (cell body) or 800 nL (projection) of 1:3 diluted AAV5-CaMKII-ChR2-eYFP or undiluted AAV5-CaMKII-eNpHR3.0-eYFP into the dmPFC (bilaterally for NpHR). For D1/D2 MSN optogenetic experiments, we injected 1 µL of AAV5-EF1a-DIO-hChR2-eYFP into the DMS of D1-Cre/A2a-Cre mice. We implanted a 1.25 mm ceramic ferrule with 200 µm fiber optic stub (Thorlabs) in either the dmPFC (cell body) or the DMS (projection and D1/D2 MSNs). For NpHR surgeries, we inserted two fiber optic cannulas bilaterally and waited either 4-6 weeks (cell body) or 6-9 weeks (projection) for expression. See **Supplement 1** for detailed injection methods and coordinates.

### Elevated Zero Maze

Behavior was recorded in a sound-dampened chamber using Ethovision XT software (Noldus). Mice were placed in a closed arm and allowed to explore the EZM for 15 minutes (fiber photometry) or 25 minutes (optogenetic manipulations).

### Fiber Photometry Recording

*In vivo* calcium data were acquired using a custom-built rig based on a previously described setup (51). Raw photoreceiver data was extracted and signals were demodulated, normalized, and peak detected. We analyzed neural activity surrounding transitions with both a 1 cm distance threshold and a 2-second time threshold. We generated peri-event time histograms (40-second window) by time-locking the neural activity (ΔF/F) to the transitions, and z-scored the ΔF/F values to the mean and standard deviation from the baseline period (−20 to −10 seconds) for each transition and averaged across animals. We then quantified the change in calcium signal from the baseline period (pre) to the 10 seconds following the transition (post). We created spatial heatmaps by dividing the EZM into sections, calculating the mean signal (ΔF/F) for each section, and normalizing from 0 to 1 for each animal. See **Supplemental Methods** for velocity thresholding.

### Optogenetic Manipulations

For ChR2, a 473 nm laser (Shanghai Laser & Optics Century Co. LTD) was used to stimulate dmPFC cell bodies (1 mW, 10 Hz, 5 ms pulse width), projection fibers in the DMS (0.5-1 mW, 10 Hz, 5 ms pulse width), and D1/D2 MSNs (continuous 200-300 µW). For NpHR, a 532 nm laser (Shanghai Laser & Optics Century Co. LTD) was used to inhibit dmPFC cell bodies and projection fibers in the DMS bilaterally (continuous 5 mW). Each trial of dmPFC cell body and projection manipulations consisted of a 5-minute baseline laser-off period followed by ten 2-minute alternating laser on/off epochs. For dmPFC-DMS projection inhibition, a 5-minute baseline was followed by four 5-minute alternating laser on/off epochs.

### Slice Electrophysiology

We injected D1-tmt mice with AAV-CaMKII-ChR2-eYFP in the dmPFC as described above. Four to six weeks after surgery, animals were terminally anesthetized and transcardially perfused, and the brain was dissected, sliced, and stored according to the protocol described in McGregor et al. (52). The DMS was identified at low power, and the area of greatest terminal field ChR2-eYFP expression was chosen for whole-cell recordings. The presence or absence of tdTomato fluorescence was used to determine if an individual cell was a D1 or D2 MSN. We excluded neurons with physiological features of interneurons (membrane tau decay <1 msec) and patched D1 and D2 neurons in randomized serial pairs. All whole-cell recordings of excitatory postsynaptic currents (EPSCs) were acquired using the setup described in McGregor et al. (52). The internal chloride concentration was calibrated such that the reversal potential of GABA_A_-mediated (disynaptic) IPSCs was −70 mV (thus, currents recorded at −70 mV were predominantly glutamatergic in origin). Experiments were performed in picrotoxin to pharmacologically isolate EPSCs. mPFC-derived EPSCs were measured at −70 mV holding potential and evoked using brief (3 msec) full-field blue (473 nm) light pulses. EPSC amplitude was defined as the average difference between the baseline holding current (0-100 msec prior to the light pulse) and the peak of the evoked EPSC, averaged over at least five trials (20-second inter-trial interval).

### Statistical Analysis

Prism 7 software (Graphpad) was used for unpaired t-test, paired t-test, Wilcoxon signed-rank test, one-way ANOVA with Tukey’s correction for multiple comparisons, and two-way repeated measures ANOVA with Sidak’s correction for multiple comparisons.

## RESULTS

### dmPFC pyramidal neurons exhibit task-related neural activity in the EZM

We first characterized the neural activity of undefined dmPFC pyramidal neurons (henceforth referred to as “whole population dmPFC”) during approach-avoidance behavior. We virally expressed CaMKII-GCaMP6f and implanted an optical fiber (400 µm) in the dmPFC. We then recorded bulk Ca^2+^ fluorescence changes during exploration of the EZM (**Figure 1A**). To visualize neural activity spatially, we subdivided the maze into sections and calculated the mean Ca^2+^ signal in each section. We used four sections for each half of the open and closed arms; section 1 was closest to the open/closed transition point, while section 4 was in the middle of the arm (**Figure 1B**). The Ca^2+^ signal from dmPFC pyramidal neurons was lowest when mice were in the middle of a closed arm (C4), and it increased as mice approached an open arm, with the highest signal occurring in the middle of the open arm (O4) (**Figure 1B**).

**Figure 1.**
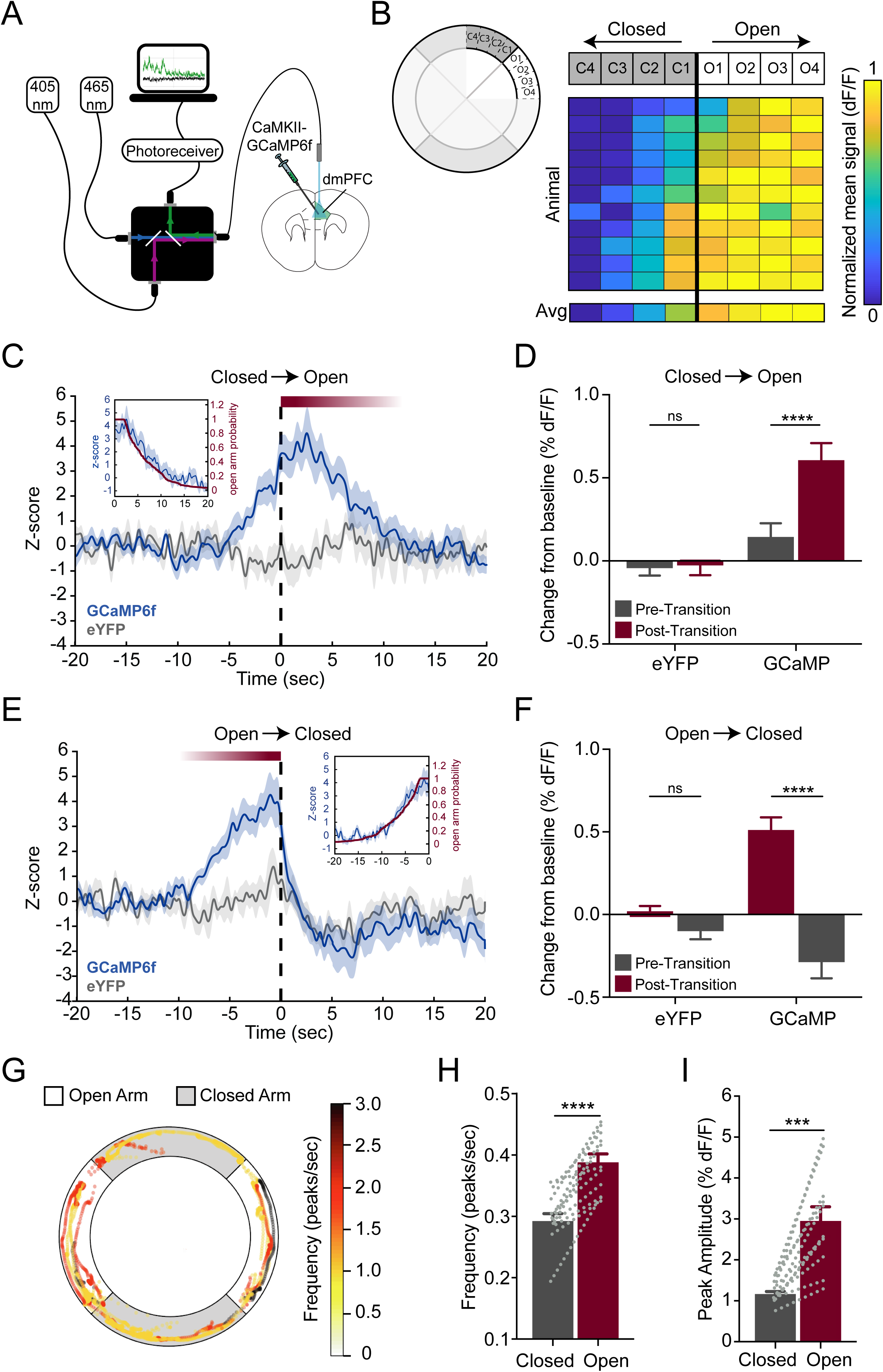
Dorsomedial prefrontal cortex pyramidal neurons exhibit task-related neural activity in the elevated zero maze. **(A)** Experimental design for Ca^2+^ imaging of dmPFC pyramidal neurons. CaMKII-GCaMP6f was virally expressed, a 400 µm optical fiber was implanted in the dorsomedial prefrontal cortex (dmPFC), and Ca^2+^ signals were recorded during exploration of the elevated zero maze (EZM). **(B)** Left, schematic of EZM with spatial sectioning. Right, mice show increased Ca^2+^ signal as they transition from the closed arms to the open arms (black line = transition point). Individual animal data is sorted by average signal across the closed arms (low to high). Average (avg) heatmap is plotted below (N_mice_ = 11). **(C)** Peri-event time histogram shows increased Ca^2+^ signal upon transition from the closed to open arms (transition at time = 0, dotted black line). The open arm GCaMP signal following the transition tightly follows the decay slope of the probability of the mice being in the open arms (inset, red line). Signal is plotted as z-score, normalized using the mean and standard deviation of the baseline period (−20 to −10 seconds) (N_transitions, GCaMP_ = 204, N_transitions, eYFP_ = 127, blue line = mean +/- SEM GCaMP, gray line = mean +/- SEM eYFP control signal, gray shading = SEM eYFP control signal). **(D)** Ca^2+^ signal is significantly higher in the open arm post-transition period (0 to 10 seconds) than in the closed arm pre-transition period (−10 to 0 seconds). No significant difference is seen in the eYFP control signal. **(E)** Same as (C) for the open to closed arm transition. Mice show a decrease in signal following transition into the closed arms. The open arm GCaMP signal preceding the transition tightly follows the decay slope of the probability of the mice being in the open arms (N_transitions, GCaMP_ = 226, N_transitions, eYFP_ = 150). **(F)** Ca^2+^ signal (% ΔF/F normalized to baseline ΔF/F; baseline is −20 to −10 seconds) is significantly lower in the closed arm post-transition period (0 to 10 seconds) than in the open arm pre-transition period (−10 to 0 seconds). No significant difference is seen in the eYFP control signal. **(G)** Aerial view of the EZM shows a representative individual animal’s trajectory (in 5-second bins), color-coded by frequency of Ca^2+^ transients with higher frequency in the open arms than in the closed arms. **(H)** Frequency of Ca^2+^ transients is higher in the open arms than in the closed arms. **(I)** Peak amplitude of Ca^2+^ transients is higher in the open arms than in the closed arms.

We also examined temporal changes in the neural signal surrounding the open/closed arm transitions. We plotted a peri-event time histogram (PETH) of the Ca^2+^ signal for the +/- 20 seconds surrounding each transition (closed-to-open and open-to-closed). Average Ca^2+^ signal was generated for three different time windows: baseline (−20 to −10 seconds), pre-transition (−10 to 0 seconds), and post-transition (0 to 10 seconds). dmPFC neurons showed a significant increase in signal as mice transitioned from closed to open arms (**Figures 1C and 1D**, two-way RM ANOVA interaction, F(_1,329_) = 17.7, p < 0.0001; Sidak’s multiple comparisons, p < 0.0001 (GCaMP, pre-transition vs. post-transition), p = 0.9727 (eYFP, pre-transition vs. post-transition); N_transitions, GCaMP_ = 204, N_transitions, eYFP_ = 127). Paralleling the spatial heatmap findings, the increase in Ca^2+^ signal slightly preceded the transition into the open arms. Conversely, dmPFC neurons showed a significant decrease in signal as animals transitioned from open to closed arms (**Figures 1E and 1F**, two-way RM ANOVA interaction, F_(1,374)_ = 44.25, p < 0.0001; Sidak’s multiple comparisons, p < 0.0001 (GCaMP, pre-transition vs. post-transition) p = 0.3962 (eYFP, pre-transition vs. post-transition); N_transitions, GCaMP_ = 204, N_transitions, eYFP_ = 127). Unlike the gradual change in signal seen in the closed-to-open transition, the signal decayed rapidly upon return to the closed arms. Control eYFP animals showed no signal modulation during either transition. We plotted the probability of the mice being in the open arms at any given timepoint (**Figure 1C, E** inset); the decay slope in the Ca^2+^ signal tightly parallels the probability that the mouse is in the open arms, and the decay duration matches the average time spent in the open arms. Together, these spatiotemporal changes indicate that on average, dmPFC activity increases as the mice approach and enter an open arm and then decreases as they transition back into a closed arm.

In addition to quantifying changes in neural activity surrounding the transition zone, we compared additional measures of neural activity between the open and closed arms. To visualize the frequency of Ca^2+^ events, we calculated frequency of event peaks in 5-second bins and plotted frequency as a function of spatial location in the EZM (**Figure 1G**). dmPFC pyramidal neurons showed a higher frequency of Ca^2+^ events in the open arms than in the closed arms (**Figure 1H**, paired t-test, p < 0.0001; N_mice_ = 11). dmPFC pyramidal neurons also showed significantly greater average peak amplitude of Ca^2+^ events in the open arms than in the closed arms (**Figure 1I**, paired t-test, p = 0.0002; N_mice_ = 11). To control for any differences in velocity of movement in the open versus closed arms, we analyzed the neural signal during bouts of similar velocity in the closed and open arms (any bout during which the animal moved < 7cm/sec for 10 seconds or longer was discarded). Using this velocity thresholding, we found that these open-arm-related changes in neural activity did not depend on velocity (**Figure S1**). Taken together, these results indicate that the activity of dmPFC pyramidal neurons is lowest in the closed arms, increases as mice approach the open arms, and peaks in the open arms, suggesting that these neurons are encoding aspects of this approach-avoidance task.

### Fronto-striatal, but not fronto-amygdalar, projection neurons recapitulate whole population dmPFC activity in the EZM

Whole population recording does not provide projection-specific information about dmPFC neurons involved in approach-avoidance behavior and may mask the activity of less-represented subpopulations in the dmPFC. We therefore next recorded the activity of projection-defined subpopulations of dmPFC neurons during exploration of the EZM. While the dmPFC-BLA projection has been well-studied in fear expression, a recent study showed that this projection is not causally involved in approach-avoidance behavior (29). We thus hypothesized that a different subpopulation of dmPFC neurons – the fronto-striatal projection to the DMS – drives the encoding of approach-avoidance behavior we observed at the whole population level. Recently, the DMS was found to have a causal role in approach-avoidance behavior in the EZM (44), and optogenetic manipulation of the dmPFC-DMS projection causally modulates decision-making under conflict, a prefrontal function relevant to approach-avoidance conflict (49).

To examine the roles of the fronto-striatal and fronto-amygdalar projections, we used a retrograde viral targeting strategy to express GCaMP6f selectively in cells projecting to either the BLA or DMS (**Figures 2A and 2E**). We injected a retrograde canine adenovirus CAV2 carrying Cre recombinase (CAV2-Cre) in the downstream area to allow for expression of Cre in any neurons projecting to that area. Additionally, we injected a Cre-dependent GCaMP6f in the upstream dmPFC, which allowed for projection-specific Ca^2+^ imaging through an implanted optical fiber (400 µm) in the dmPFC. Similar to previous analyses, we first plotted the spatial modulation of neural activity in each projection. In the dmPFC-BLA projection population, we found a mixture of responses: about half of the mice showed lower activity in the closed arms, and half showed no difference or the opposite trend. The findings were inconsistent across animals; the average spatial heatmap did not show a robust increased signal, as we observed with whole population recording, as the mice moved further into the open arms (**Figure 2B**). Likewise, dmPFC-BLA projection neurons did not show any significant difference in frequency of Ca^2+^ transients (**Figure 2C**, paired t-test, p = 0.5503; N_mice_ = 9) or amplitude of Ca^2+^ transient peaks (**Figure 2D**, Wilcoxon signed-rank test, p = 0.9102; N_mice_ = 9) in open versus closed arms. Conversely, spatial activity of the dmPFC-DMS projection resembled that of the dmPFC population as a whole (**Figure 2F**), and dmPFC-DMS projection neurons showed higher frequency (**Figure 2G**, paired t-test, p = 0.0393; N_mice_ = 10) and amplitude (**Figure 2H**, paired t-test, p = 0.0067; N_mice_ = 10) of Ca^2+^ transients in the open arms than in the closed arms. When comparing epochs of equal movement velocity between open and closed arms, dmPFC-DMS projection neurons still showed a significantly higher peak amplitude in the open arms (**Figure S1**), whereas the dmPFC-BLA peak amplitude remained unchanged across arms. These data suggest that the dmPFC-DMS population more robustly represents aspects of the approach-avoidance task than the dmPFC-BLA projection.

**Figure 2.**
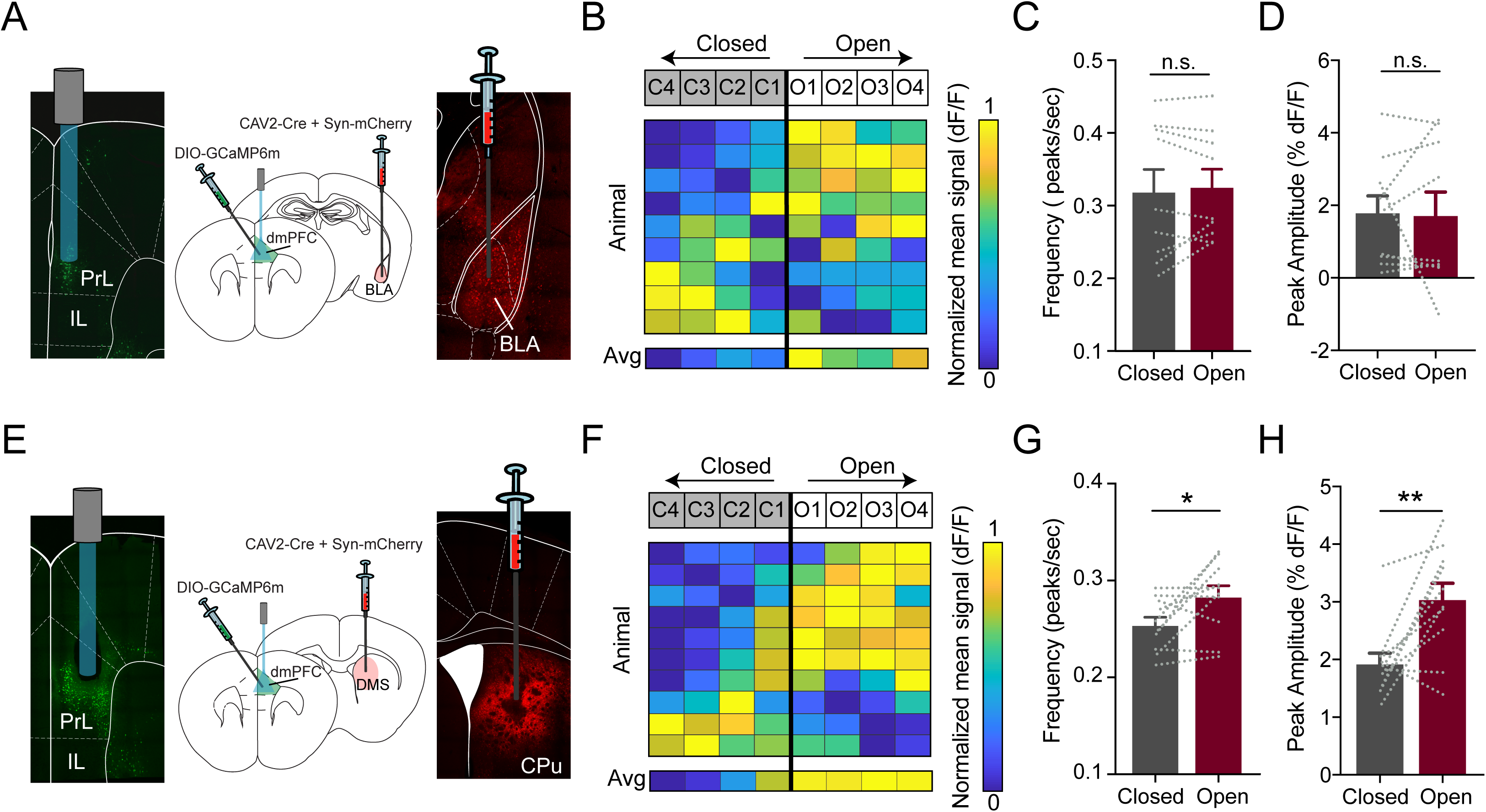
Fronto-striatal but not fronto-amygdala projection neuron activity recapitulates whole population prefrontal activity in the elevated zero maze. **(A)** Injection schematic with representative histology images shows targeting of DIO-GCaMP6m to the dorsomedial prefrontal cortex (dmPFC) and CAV-Cre / Syn-mCherry to the basolateral amygdala (BLA). **(B)** Mice show a mixture of Ca^2+^ signal values as they transition from the closed arms to the open arms (black line = transition point). Individual animal data is sorted by average signal across the closed arms (low to high). Average (avg) heatmap is plotted below (N_mice_ = 9). **(C)** Frequency of Ca^2+^ transients is similar in the open and closed arms. **(D)** Peak amplitude of Ca^2+^ transients is similar in the open and closed arms. **(E)** Injection schematic with representative histology images shows targeting of DIO-GCaMP6m to the dmPFC and CAV-Cre / Syn-mCherry to the dorsomedial striatum. **(F)** Mice show increased Ca^2+^ signal as they transition from the closed arms to the open arms (black line = transition point); the lowest and highest signals occur in the center of the closed and open arms, respectively. Individual animal data is sorted by average signal across the closed arms (low to high). Average (avg) heatmap is plotted below (N_mice_ = 10). **(G)** Frequency of Ca^2+^ transients is higher in the open arms than in the closed arms. **(H)** Peak amplitude of Ca^2+^ transients is higher in the open arms than in the closed arms.

### Optogenetic stimulation of dmPFC-DMS projection terminals preferentially excites postsynaptic D1 MSNs

The results above indicate a role for the dmPFC-DMS projection in approach-avoidance behavior, so we next investigated dmPFC-DMS connectivity. Using patch clamp electrophysiology in striatal slices combined with terminal field optogenetic stimulation of dmPFC inputs, we assessed the responses of DMS D1 and D2 MSNs to excitation of dmPFC inputs. Sequential pairs of nearby D1 and D2 MSNs were patched in the whole-cell configuration (**Figure 3A**), and both showed EPSCs in response to blue light stimulation (**Figure 3B**). We plotted the ratio of EPSCs for each recorded pair (D1 and D2) (**Figure 3C**); in almost all pairs, we observed larger EPSCs in D1 MSNs (**Figure 3D**, Wilcoxon signed-rank test, p = 0.0054; N_cell pairs_ = 14), yielding a ratio > 1. These results indicate that the dmPFC projection preferentially activates D1 MSNs.

**Figure 3.**
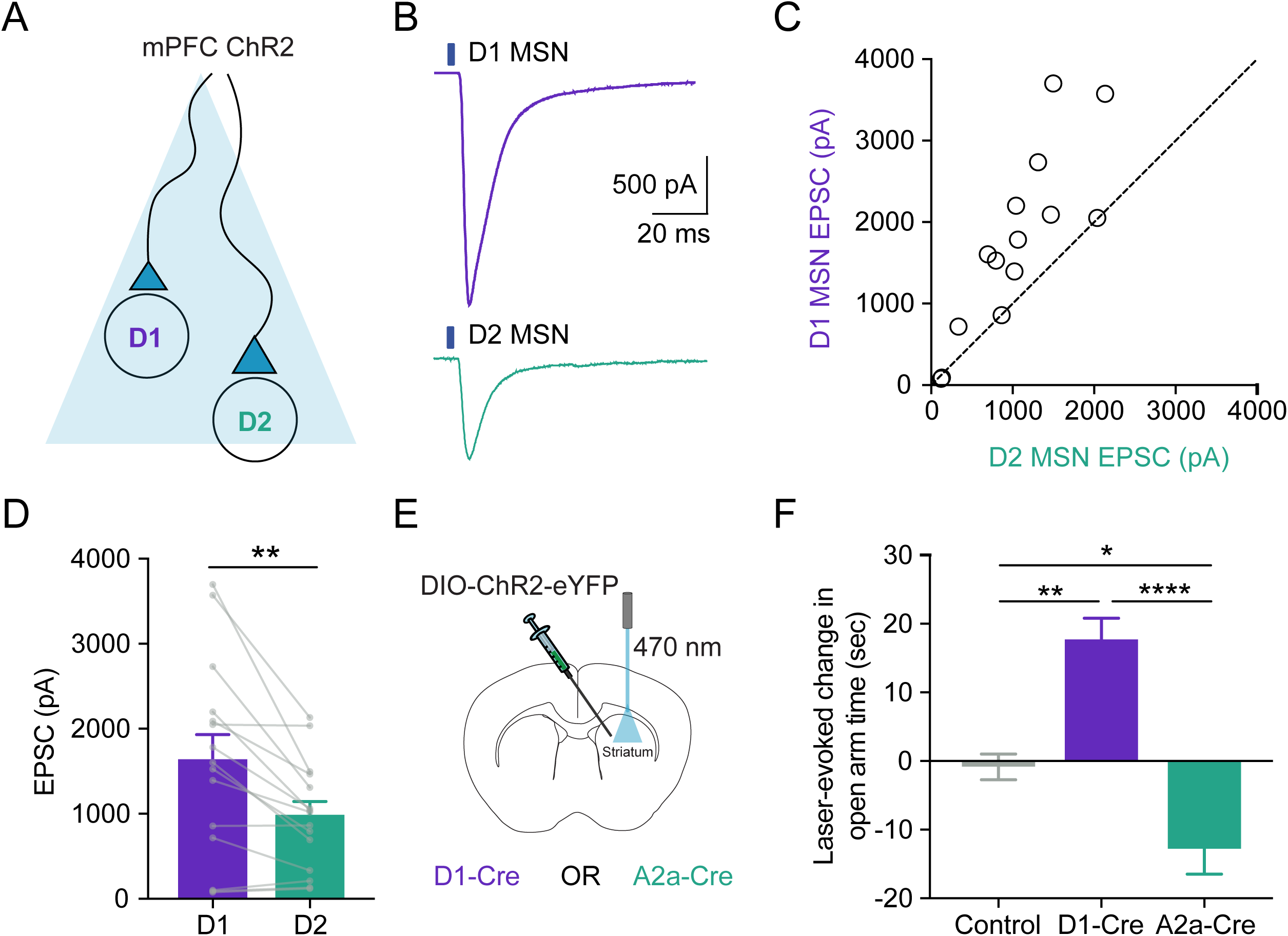
Prefrontal projections preferentially recruit striatal D1 medium spiny neurons that directly control avoidance behavior. **(A)** Slice electrophysiology experimental design for optogenetic stimulation of dorsomedial prefrontal cortex (dmPFC) terminals in the dorsomedial striatum (DMS). CaMKII-ChR2-eYFP was expressed in the dmPFC of mice and slice electrophysiology recordings were taken in the DMS during blue light stimulation (470 nm). **(B)** Representative traces from D1 and D2 medium spiny neurons (MSNs) in the DMS. D1 MSNs have larger excitatory postsynaptic current (EPSC) following blue light stimulation. **(C)** Paired-sequential recordings from D1 and D2 MSNs show preferential excitation of D1 MSNs following stimulation of dmPFC-DMS projection terminals. **(D)** D1 MSNs show a higher-amplitude EPSC than D2 MSNs in response to stimulation. **(E)** Schematic of optogenetic stimulation of D1 or D2 MSNs *in vivo*. DIO-ChR2-eYFP was virally expressed in the DMS of either D1-Cre +/- or A2a Cre +/- mice. A 200 µm optical fiber was implanted in the DMS, and mice were optogenetically-stimulated (via 470 nm light) during exploration of the EZM. **(F)** Optogenetic stimulation of D1 MSNs increased the time spent in the open arms, while stimulation of D2 MSNs decreased the time spent in the open arms. Control animals showed no effect of laser stimulation.

### Optogenetic stimulation of D1 and D2 MSNs in the DMS bidirectionally modulates approach-avoidance behavior

Given the stronger excitation of D1 than D2 MSNs by dmPFC inputs, we next tested the causal effects of directly stimulating D1 and D2 MSNs on approach-avoidance behavior. Using D1-Cre and A2a-Cre mouse lines to target these two populations, respectively, we expressed Channelrhodopsin-2 (ChR2) in either D1 or D2 MSNs by injecting a Cre-dependent ChR2 virus into the DMS. We then implanted a 200 µm optical fiber above the area of expression to allow for direct stimulation of D1 and D2 MSNs during exploration of the EZM (**Figure 3E**). We followed a 25-minute paradigm consisting of 5 minutes of baseline exploration followed by 2-minute alternating epochs of laser on/off. D1-Cre mice spent significantly more time exploring the open arms in laser-on epochs than in laser-off epochs, whereas A2a-Cre mice spent significantly less time exploring the open arms when the laser was on (**Figure 3F**, one-way ANOVA interaction, F_(2,23)_ = 22.39, p < 0.0001; Tukey’s multiple comparisons, p = 0.0013 (Control vs. D1-Cre), p = 0.0160 (Control vs. D2-Cre), p < 0.0001 (D1-Cre vs. D2-Cre); N_D1-Cre_ = 6 mice, N_A2a-Cre_ = 10 mice, N_eYFP_ = 10 mice). We did not observe any laser-evoked differences in open arm time in control animals, which were a mix of D1-Cre/A2a-Cre mice expressing Cre-dependent eYFP and wild-type littermates with DMS optical fiber implants to control for the effects of viral expression, surgery, laser, and optical patchcord tethering. Light stimulation had no significant effect on locomotion in the control or D1-Cre groups but decreased locomotion in the A2a-Cre group (**Figure S2**). Following optogenetic stimulation of D1 MSNs, we next investigated the effect of inhibiting these neurons via chemogenetic inactivation using inhibitory designer receptors exclusively activated by designer drugs (DREADDs). We injected a Cre-dependent hM4Di virus in the DMS of D1-Cre mice and injected clozapine-N-oxide (CNO) 10 minutes prior to exploration of the elevated plus maze (EPM). Chemogenetic inactivation of D1 MSNs decreased the time spent in the open arms but did not affect locomotion (**Figure S3**). Together, these data indicate that D1 MSN activity drives exploratory approach behavior.

### Optogenetic manipulation of dmPFC-DMS projection neurons bidirectionally controls approach-avoidance behavior

Given our findings that endogenous activity of dmPFC-DMS projection neurons was highest in the open arms, and that stimulating DMS D1 MSNs causally increased open arm exploration, we hypothesized that dmPFC inputs may provide a necessary source of excitation to drive exploratory behavior via downstream D1 MSNs. We therefore tested whether using optogenetic manipulation to increase the activity of dmPFC-DMS projections, augmenting the increase in activity naturally observed in the open arms, would increase open arm exploration. To this end, we expressed CaMKII-ChR2-eYFP in the dmPFC of mice and implanted an optical fiber (200 µm) in the DMS to allow for *in vivo* optogenetic stimulation of dmPFC-DMS terminals during exploration of the EZM (**Figure 4A**). ChR2 mice spent significantly more time exploring the open arms in laser-on epochs compared with laser-off epochs, and there was no effect of laser in eYFP controls (**Figure 4B-D**, two-way RM ANOVA interaction, F_1,15_ = 14.03, p = 0.0019; Sidak’s multiple comparisons, p < 0.0001 (ChR2, laser on vs. off), p = 0.9256 (eYFP, laser on vs. off); N_ChR2_ = 9 mice, N_eYFP_ = 8 mice). Additionally, ChR2 mice spent significantly more time in the open arms during the last 5 minutes than during the baseline period (pre-stimulation), while control animals showed no difference (**Figure 4E**, two-way RM ANOVA interaction, F_1,15_, p = 0.0017; Sidak’s multiple comparisons, p = 0.0084 (ChR2, first 5 minutes vs. last 5 minutes), p = 0.1142 (eYFP, first 5 minutes vs. last 5 minutes); N_ChR2_ = 9 mice, N_eYFP_ = 8 mice). Laser stimulation had no effect on locomotion in either of the groups (**Figure S4A**). While optogenetic stimulation of dmPFC-DMS terminals was sufficient to increase approach behavior in the EZM, we next tested whether activity in this pathway is necessary for normal approach-avoidance behavior. We expressed CaMKII-eNpHR3.0-eYFP in the dmPFC of mice and implanted a 200 µm optical fiber in the downstream DMS to allow for optogenetic inhibition of projection terminals during exploration of the EZM (**Figure S4B**). Optogenetic inhibition of these terminals significantly decreased time spent in the open arms during the laser-on epochs relative to the laser-off epochs (**Figure S4C**). There was no effect of laser on locomotion or on behavior in control eYFP mice (**Figure S4D**). Finally, we tested the effect of non-projection-specific whole population dmPFC pyramidal neuron optogenetic activation/inhibition and found no significant effect on approach-avoidance behavior, suggesting that our behavioral effects were specific to the fronto-striatal projection (**Figure S5**). These data suggest that activation of the dmPFC-DMS pathway is both necessary and sufficient for approach-avoidance behavior in the EZM.

**Figure 4.**
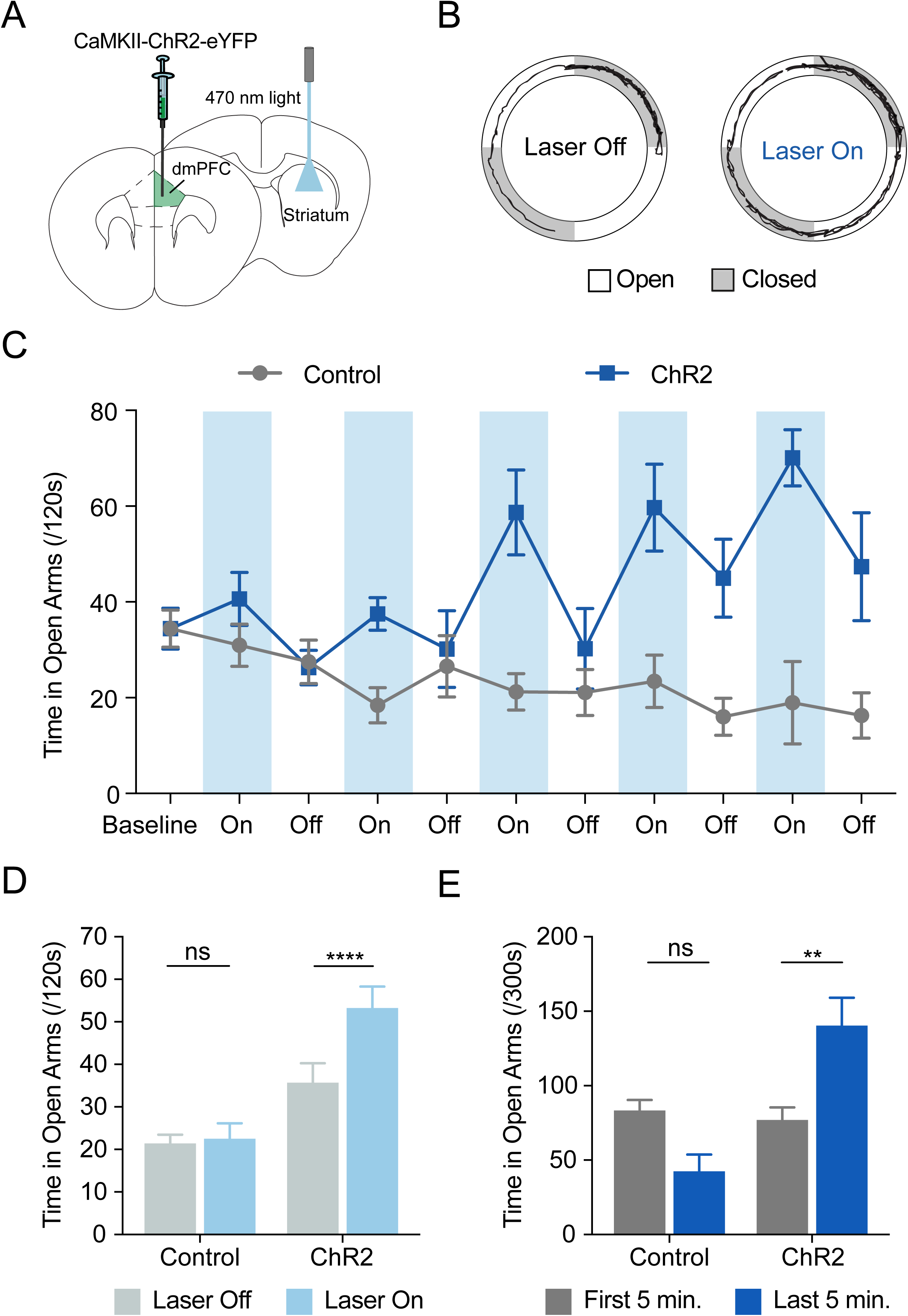
Optogenetic stimulation of fronto-striatal neurons decreases avoidance behavior in the elevated zero maze. **(A)** Schematic showing optogenetic stimulation of dorsomedial prefrontal cortex (dmPFC) projections to the dorsomedial striatum. CaMKII-ChR2-eYFP was virally expressed in the dmPFC, and a 200 µm optical fiber was implanted in the DMS. Mice were optogenetically stimulated (470 nm light) during exploration of the elevated zero maze. **(B)** Representative individual animal trace shows increased exploration of the open arms during stimulation (laser on). **(C)** ChR2 mice show a selective increase in time spent in open arms during laser-on epochs across the entire stimulation paradigm (5-minute baseline followed by 2-minute on/off epoch of laser stimulation). Control mice show no modulation of time spent in open arms in response to laser stimulation. **(D)** ChR2 mice show an increase in open arm time during laser-on epochs than in laser-off epochs. Control eYFP mice show no modulation of open arm time in laser-on versus laser-off epochs. **(E)** ChR2 mice show an increase in open arm time during the last 5 minutes of exploration (following stimulation) than in the first 5 minutes (baseline, pre-stimulation). Control eYFP mice trend toward less open arm time in the last 5 minutes of exploration than in the first 5 minutes.

## DISCUSSION

We found that dmPFC pyramidal neurons on average exhibit an increase in activity during approach and exploration of the open arms of the EZM, corroborating previous studies showing that mPFC units distinguish between the open and closed arms (28). The mPFC is well-situated to play a critical role in processing approach-avoidance conflict, as it receives inputs carrying contextual and valence information. Specifically, inputs from the BLA and ventral hippocampus (vHPC) to the mPFC are required for normal expression of approach-avoidance behavior (53, 54). However, little previous work has compared the roles of distinct efferent projections of the dmPFC in approach-avoidance behavior. Here, we addressed this knowledge gap by investigating differential representation of approach-avoidance behavior by fronto-striatal and fronto-amygdala projection neurons. Ca^2+^ signals from dmPFC-BLA projection neurons did not indicate substantial differences in neural activity during exploration of open and closed arms of the EZM. This result was particularly striking given that previous studies have focused on the dmPFC-BLA projection for its role in controlling fear expression (24, 55-57) and have implicated the mPFC-BLA projection in safety signaling (58, 59). In the context of this previous data, our results suggest that while the dmPFC-BLA projection may be important for reflexive defensive behaviors such as freezing, a different top-down dmPFC projection may be involved in the more cognitively demanding, approach-avoidance decision-making behaviors relevant to anxiety. This model is supported by a recent study in which optogenetic stimulation of the dmPFC-BLA projection did not affect approach-avoidance behavior but did affect cued freezing during fear extinction retrieval (29).

We then turned to an alternative dmPFC projection target, the DMS, which is implicated in controlling action selection (43), goal-directed actions (60), and more recently approach-avoidance behavior (44). The DMS is well situated to receive action-initiation or inhibition signals from the dmPFC to facilitate approach or avoidance movements through its projection to downstream basal ganglia targets. Of particular relevance is a recent study that investigated the role of the dmPFC-DMS projection in an alternative approach-avoidance conflict task. This experiment used a modified T-maze with a mixture of explicit costs and rewards (bright light bulbs and chocolate milk, respectively) at the end of each arm. One condition provided a “cost-benefit conflict”—one arm contained concentrated chocolate milk and a bright light (high-cost, high-reward), and the other arm contained dilute chocolate milk and a dim light (lost-cost, low-reward)—while other conditions allowed for choices without conflict (e.g. bright light vs. dim light). dmPFC-DMS projection neurons robustly increased activity at the decision point of the T-maze *only* under conflict conditions, suggesting that the dmPFC-DMS projection is active during decision-making in approach-avoidance conflict situations but not in general value-based decision-making (49). In alignment with this previous work, we found that dmPFC-DMS projection neurons encoded aspects of our EZM approach-avoidance task, with greater activity in the open arms than in the closed arms. Given the increase in neural activity preceding entrance into the open arms, the dmPFC-DMS projection neurons may be responsive to the decision occurring at the transition point. In this “risk assessment” zone, an increase in neural activity would drive approach toward the open arms. These findings, combined with the previous work, suggest a model in which distinct subpopulations of dmPFC projection neurons play differential roles in anxiety-related behaviors, with the dmPFC-BLA projection involved primarily in reflexive fear behavior and the dmPFC-DMS projection involved primarily in cognitive anxious avoidance behavior.

Although our fiber photometry results show increased activity of dmPFC-DMS projection neurons during exploration of the open arms, it is not possible to interpret the directionality or valence of this signal from Ca^2+^ imaging alone. Theoretically, this increased signal could be interpreted in two opposing ways: as a correlate of increased “anxiety” that the animals experience after entering the open arms, or as a correlate of decreased “anxiety” that drove the animals into the open arms. In order to discriminate between these two possibilities and causally link the dmPFC-DMS projection to approach-avoidance behavior, we employed projection-specific optogenetic stimulation in the EZM. We found that fronto-striatal projection stimulation increased open arm exploration, while inhibition decreased open arm exploration. These results, combined with our Ca^2+^ imaging data, suggest that the increase in endogenous fronto-striatal activity in the open arms is likely a correlate of increased approach behavior or decreased anxiety-like behavior. This result is surprising given the classical role of the dmPFC in fear conditioning, in which increased dmPFC activity is associated with increased fear expression (19-21). When combined with our finding that global optogenetic manipulation of the dmPFC as a whole has no effect on approach-avoidance behavior, this result highlights the importance of considering projection specificity when addressing the heterogeneous dmPFC.

After identifying this novel role for dmPFC-DMS projections in encoding and controlling approach-avoidance behavior, we further characterized the dmPFC-DMS circuit at the synaptic level and investigated the role of different downstream cell types within the DMS. One previous rabies tracing study suggested that the dmPFC preferentially innervates striatal D1 MSNs (61), while another study found similar innervation of D1 and D2 MSNs (62). Using slice physiology, we confirmed that stimulation of dmPFC projection fibers in the DMS preferentially activated D1 MSNs, although we also found appreciable activation of D2 MSNs. We then looked more closely at the behavioral relevance of different circuitry within the striatum itself. D1 and D2 MSNs have been shown to play opposing roles in reward and reinforcement; specifically, stimulation of D1 MSNs is reinforcing, while stimulation of D2 MSNs is punishing (42, 63, 64). A recent study found that optogenetic stimulation of D2 MSNs increased avoidance behavior in the EZM, while chemogenetic inhibition of D2 MSNs reduced avoidance behavior (44). In line with and building upon this previous work, we found that stimulation of D1 MSNs decreased avoidance while stimulation of D2 MSNs increased avoidance in the EZM. These results indicate that in addition to playing opposing roles in movement and reinforcement, D1 and D2 MSNs also play opposing roles in the control of approach-avoidance behavior.

While previous studies have implicated the dmPFC and DMS separately in approach-avoidance behavior, and have implicated the dmPFC-DMS circuit in decision-making under conflict (49), our findings build upon this previous work by providing direct evidence that the dmPFC-DMS is a novel circuit involved in controlling anxiety-like behavior in the EZM, while the dmPFC-BLA pathway does not play a robust role. Our results support a model for prefrontal control of defensive behavior in which fronto-striatal projection neurons modulate defensive *actions* such as avoidance, and fronto-amygdalar projection neurons modulate defensive *reactions* such as freezing. This model may be solidified by further studies during fear behaviors to demonstrate selective recruitment of the dmPFC-BLA projection, not the dmPFC-DMS projection. Additionally, it is not known whether the dmPFC-DMS circuit is specific to innate approach-avoidance behavior or more broadly involved in other types of learned avoidance behavior, such as active and passive avoidance. As such, future studies should compare the neural representations of these different types of avoidance behavior in a circuit-specific manner.

A core feature of human anxiety disorders is excessive avoidance behavior, which presents a barrier to treating these disorders. Our findings identify a novel fronto-striatal circuit that controls approach-avoidance conflict and may be a valuable target for future animal and human studies that seek to restore balance between approach and avoidance behaviors.

## ACKOWLEDGEMENTS AND DISCLOSURES

This study was funded by a Chan-Zuckerberg Biohub Investigator Award and a Weill Innovation Award to LAG. ACK was supported by NIH R01 NS064984.

All authors report no potential conflicts of interest.

