## Supplemental Material for "Fronto-striatal projections regulate approach-avoidance conflict"

**SUPPLEMENTAL METHODS (Supplement 1)**

All experiments were conducted in accordance with procedures established by the Administrative Panels on Laboratory Animal Care at the University of California, San Francisco.

**Animal Subjects**

We used wild-type C57BL/6J (The Jackson Laboratory), Tg(Drd1a-cre)EY217Gsat (The Jackson Laboratory), Tg(Adora2a-cre)KG139Gsat (The Jackson Laboratory), and Drd1a-tdTomato (D1-tmt) mice (50). D1-Cre, A2a-Cre, and D1-tmt mice were on an C57BL/6J background. All mice were raised in normal light conditions (12:12 light/dark cycle), fed and watered *ad libitum.*

**Stereotaxic Surgery**

Surgeries were done at 10-14 weeks of age. Mice were anesthetized using 5.0% isoflurane at an oxygen flow rate of 1 L/min and placed on top of a heating pad in a stereotaxic apparatus (Kopf Instruments). Anesthesia was maintained with 1.5-2.0% isoflurane for the duration of the surgery. Respiration and pinch response were monitored closely. Slow-release buprenorphine (.5mg/kg) and ketoprophen (1.6 mg/kg) were administered subcutaneously at the start of surgery. The incision area was shaved and cleaned with ethanol and betadine. Lidocane (0.5%) was administered topically on the scalp. An incision was made along the midline and bregma was measured. Virus was injected (as described below) using a 10 µL nanofil syringe (World Precision Instruments) with a 33-gauge beveled needle. The needle was facing anterior for dmPFC injections and medial for DMS/BLA injections. We used an injection rate of 100 nL/min with a 10-minute delay before retracting the needle. Mice recovered in a clean cage on top of a heating pad and a subsequent injection of ketoprofen (1.6 mg/kg) was given the following day.

**Injection of virus for GCaMP expression (fiber photometry)**

*dmPFC pyramidal neurons:*

We expressed GCaMP6f in dmPFC pyramidal neurons using an adeno-associated virus (AAV) vector with serotype 5 to drive the expression of GCaMP6f-WPRE-SV40 under the CaMKII promoter in wild-type mice. Coordinates (in millimeters relative to bregma) for injection into the dmPFC were 1.8 A/P, -.35 M/L, -2.6 D/V. We injected 500 nL of virus and waited 4-5 weeks for expression before recording. For control animals we injected 500 nL of AAV5-CaMKIIa-eYFP (UNC Vector Core) into the dmPFC.

pENN.AAV.CamKII.GCaMP6f.WPRE.SV40 was a gift from James M. Wilson (Addgene viral prep # 100834-AAV5; http://n2t.net/addgene:100834 ; RRID:Addgene_100834). Titer 4.12E+13. AAV5-CaMKIIa-eYFP was a gift from Karl Deisseroth and packaged by the UNC Vector Core. Titer 3.60E+12.

*dmPFC-DMS and dmPFC-BLA projection neurons:*

We expressed GCaMP6m in dmPFC neurons projecting to either the DMS or BLA using a dual virus retrograde targeting strategy in wild-type mice. We used an adeno-associated virus (AAV) vector with serotype 1 to drive the expression of the Cre-dependent construct Flex-GCaMP6m-WPRE-SV40 under the synapsin (Syn) promoter (Addgene) in the dmPFC. Additionally, in the downstream target region (DMS or BLA) we injected a retrograde canine adenovirus type 2 (CAV2) to drive expression of Cre recombinase (Institut de Génétique Moléculaire de Montpellier, Montpellier, France) as well as an AAV8 virus to drive the expression of mCherry under the human synapsin (hSyn) promotor (UNC Vector Core; to visualize CAV2 injection location). Coordinates (in millimeters relative to bregma) for dmPFC injections were the same as above, coordinates for the DMS were .8 A/P, -1.5 M/L, -3.5 D/V, and coordinates for the BLA were -1.4 A/P, -3.3 M/L, -4.9 D/V. We injected 1500 nL of Syn-Flex-GCaMP6m in the dmPFC and either 350 nL of each CAV2-Cre and hSyn-mCherry in the DMS or 250 nL of each in the BLA. We waited 4-5 weeks for expression before recording. For control animals we injected 1500 nL AAV5-EF1a-DIO-eYFP-WPRE-hGH (Addgene) into the dmPFC.

pAAV.Syn.Flex.GCaMP6m.WPRE.SV40 was a gift from The Genetically Encoded Neuronal Indicator and Effector Project (GENIE) & Douglas Kim (Addgene viral prep # 100838-AAV1; http://n2t.net/addgene:100838 ; RRID:Addgene_100838). Titer 2.10E+13. pAAV-Ef1a-DIO EYFP was a gift from Karl Deisseroth (Addgene viral prep # 27056-AAV5 ; http://n2t.net/addgene:27056 ; RRID:Addgene_27056). Titer 2.40E+13. CAV2-Cre was packaged by the Plateforme de Vectorologie de Montpellier. Titer 1.00E+13. AAV8-hSyn-mCherry was a gift from Karl Deisseroth and packaged by the UNC Vector Core. Titer 4.1E +12.

**Injection of virus for optogenetic manipulations**

All injections were unilateral (left hemisphere) unless otherwise specified.

*dmPFC pyramidal neurons:*

We expressed channelrhodopsin (ChR2) or halorhodopsin (NpHR) in dmPFC pyramidal neurons using an adeno-associated virus (AAV) vector with serotype 5 to drive the expression of hChR2-eYFP-WPRE-hGH (Addgene) or eNpHR3.0-eYFP (UNC Vector Core) under the control of a CaMKIIa promoter in wild-type mice. Coordinates (in millimeters relative to bregma) for injection into the dmPFC were 1.8 A/P, -0.35 M/L, -2.6D/V. We injected 500 nL of virus (1:3 dilution in saline for ChR2, undiluted for NpHR) at and waited 4-6 weeks for expression before running behavioral assays. For control animals we injected 500 nL of undiluted AAV5-CaMKIIa-eYFP (UNC Vector Core) into the dmPFC.

pAAV-CaMKIIa-hChR2(H134R)-EYFP was a gift from Karl Deisseroth (Addgene viral prep # 26969-AAV5; http://n2t.net/addgene:26969 ; RRID:Addgene_26969). Titer 6.10E+12. AAV5-CaMKIIa-eNpHR3.0-eYFP was a gift from Karl Deisseroth and packaged by the UNC Vector Core. Titer 4.30E+12. AAV5-CaMKIIa-eYFP was a gift from Karl Deisseroth and packaged by the UNC Vector Core. Titer 3.60E+12.

*dmPFC-DMS projection neurons:*

We expressed channelrhodopsin (ChR2) or halorhodopsin (NpHR) in dmPFC pyramidal neurons using an adeno-associated virus (AAV) vector with serotype 5 to drive the expression of hChR2-eYFP-WPRE-hGH (Addgene) or eNpHR3.0-eYFP (UNC Vector Core) under the control of a CaMKIIa promoter in wild-type mice. Coordinates (in millimeters relative to bregma) for injection into the dmPFC were 1.8 A/P, -0.3 M/L, -2.6 D/V. The NpHR virus was injected bilaterally. We injected 800 nL of 1:3 diluted ChR2 virus or 500 nL of undiluted NpHR virus and waited 6-9 weeks for expression before running behavioral assays. For control animals we injected 800 (ChR2 group) or 500 (NpHR group) nL of AAV5-CaMKIIa-eYFP (UNC Vector Core) into the dmPFC.

pAAV-CaMKIIa-hChR2(H134R)-EYFP was a gift from Karl Deisseroth (Addgene viral prep # 26969-AAV5; http://n2t.net/addgene:26969 ; RRID:Addgene_26969). Titer 6.10E+12. AAV5-CaMKIIa-eNpHR3.0-eYFP was a gift from Karl Deisseroth and packaged by the UNC Vector Core. Titer 4.30E+12. AAV5-CaMKIIa-eYFP was a gift from Karl Deisseroth and packaged by the UNC Vector Core. Titer 3.60E+12.

*D1 and D2 striatal MSNs:*

We expressed channelrhodopsin (ChR2) in DMS MSNs using an adeno-associated virus (AAV) vector with serotype 5 to drive the expression of DIO-hChR2-eYFP-WPRE-hGH (Addgene) under the control of an EF1a promoter in either D1-Cre or A2a-Cre mice. Coordinates (in millimeters relative to bregma) for injection into the DMS were (0.8 A/P, -1.5 M/L, -3.5 D/V). We injected 1 µL of virus bilaterally and waited 4 weeks for expression before running behavioral assays. For control animals we injected 1 µL of AAV5.EF1a-DIO.eYFP (Addgene) into the DMS.

AAV5-EF1a-DIO-hChR2(H134R)-eYFP was a gift from Karl Deisseroth and packaged by the UNC Vector Core. Titer 3.84E+13. pAAV5-Ef1a-DIO-EYFP was a gift from Karl Deisseroth (Addgene plasmid # 27056 ; http://n2t.net/addgene:27056 ; RRID:Addgene_27056).

**Injection of virus for chemogenetic manipulations**

We expressed an inhibitory designer receptor exclusively activated by designer drugs (DREADDs, hM4D(Gi)) in the DMS using adeno-associated virus (AAV) vector with serotype 5 to drive the expression of DIO-hM4D(Gi)-mCherry (Addgene) under the control of a hSyn promotor in D1-Cre mice. Coordinates (in millimeters relative to bregma) for injection into the DMS were (0.8 A/P, -1.5 M/L, -3.5 D/V). We injected 1 µL of 1:3 diluted virus (in saline) bilaterally and waited 6 weeks for expression before running behavioral assays.

AAV5-hSyn-DIO-hM4D(Gi)-mCherry was a gift from Bryan Roth (Addgene viral prep # 44362-AAV5; http://n2t.net/addgene:44362 ; RRID:Addgene_44362). Titer 1.20E+13.

**Fiber-optic cannula implantation**

For fiber photometry surgeries a 2.5 mm metal fiber optic cannula (0.48 NA, 400 µm diameter, Doric Lenses) was implanted in the dmPFC (1.8 A/P, -.35 M/L, -2.4 D/V). For optogenetic surgeries, a 1.25 mm ceramic fiber optic cannula (0.39 NA, 200 µm diameter, Thorlabs) was implanted in either the dmPFC (1.8 A/P, -0.3 M/L, -2.3 D/V) or the DMS (0.9 A/P; -1.0 M/L; -3.0 D/V for projection fibers, 0.8 A/P; -1.5 M/L; -3.0 D/V for D1/D2 MSNs). For NpHR surgeries, two fiber optic cannulas were inserted (bilaterally, either the dmPFC or DMS). All coordinates are measured from bregma. Dental cement (Metabond) was used to secure the fiber optic cannula in place and the skin was sutured around the implant.

**Behavioral Assays**

On the experiment day the fiber optic patch cord was connected to the cannula implant using a ceramic sleeve (Thorlabs). All behavioral assays were done in an open top sound-dampened chamber. Mice were then placed in the EZM.

*Elevated Zero Maze*

The EZM was custom-made using matte white plastic for the floor and closed arm walls and clear plastic for the inner wall of the closed arms (dimensions: 55 cm diameter, 60 cm tall, 30 cm mouse floor height). Mice were initially placed in a closed arm. The EZM sessions lasted 15 minutes for fiber photometry recording experiments and 25 minutes for optogenetic manipulation experiments. Time spent in open arms and closed arms was recorded and quantified by Ethovision XT software (Noldus).

**Fiber Photometry**

*Recording*

In vivo calcium imaging data were acquired using a custom-built rig based on a previously described setup (51). This setup was controlled by an RZ5P fiber photometry processor (TDT) and Synapse software (TDT). The RZ5P/Synapse software controlled a 4 channel LED Driver (DC4100, Thorlabs) which in turn controlled two fiber-coupled LEDS: 470 nm for GCaMP stimulation and 405 nm to control for artifactual fluorescence (M470F3, M405FP1, Thorlabs). These LEDs were sinusoidally modulated at 210 Hz (470 nm) and 320 Hz (405 nm) and connected to a Fluorescence Mini Cube with 4 ports (Doric Lenses) and the combined LEF output was connected through a fiber optic patch cord (0.48 NA, 400 μm, Doric Lenses) to the cannula via a ceramic sleeve (Thorlabs). The emitted light was focused onto a Visible Femtowatt Photoreceiver Module (Model 2151, Newport, AC low) and sampled at (60 Hz).Video tracking software (Ethovision, Noldus) was time synced to the photometry setup using TTL pulses generated every 10 seconds following the start of the Noldus trial.

*Data Analysis*

Raw photoreceiver data was extracted and analyzed using custom scripts in Matlab R2018b (The MathWorks). The two output signal data was demodulated from the raw signal based on the LED modulation frequency. To normalize the data and correct for bleaching, the 405 nm channel signal was fitted to a polynomial over time and subtracted from the 470 nm GCaMP signal, giving the $\Delta F/F$ value. For detection and quantification of peak amplitude and frequency of calcium transients, we used a custom-written peak detection algorithm using a running average method, a 10-second smoothing window for determining peak to trough height and a minimum temporal resolution of 1 second between peaks. Average peak amplitude and frequency of calcium transients were compared when the animal was in the open arms as compared to the closed arms averaged across all animals. For analyzing neural activity surrounding transitions (i.e. when the animal moves from the open to closed are and vice versa), we used both a 1 cm distance threshold (i.e. the animal center point must be at least 1 cm into the closed/open arm to be considered a transition) as well as a 2-second time threshold (i.e. the animal must be in the new arm for at least 2 seconds). We time-locked the neural activity (∆F/F) to the time of these transitions as generated by Ethovision XT using a pre and post 20-second window. We then z-scored the ∆F/F values to the mean and standard deviation from the "baseline period" (defined as -20 to -10 seconds) for each transition and averaged this across animals to generate a peri-event time histogram (PETH) graph. We then quantified the average change in neural signal from baseline average signal in the 10 seconds preceding the transition (pre) as compared to the 10 seconds following the transition (post) and averaged this across animals. For creating the spatial heatmaps we divided the EZM into sections and calculated the mean signal (∆F/F) when the animal was in each of the sections and normalized from 0 to 1 for each animal. For velocity thresholding we used a threshold of 7 cm/sec for 10 seconds (i.e. any bout where the animal is under 7 cm/sec for 10 seconds or longer is discarded in comparison) to allow for comparison between similar activity epochs in the closed and open arms.

**Optogenetic Manipulations**

For ChR2 experiments, blue light was generated by a 473 nm laser (Shanghai Laser & Optics Century Co. LTD) to stimulate dmPFC cell bodies (1 mW, 10 Hz, 5 ms pulse width), projection fibers in the DMS (.5-1 mW, 10 Hz, 5 ms pulse width), and D1/D2 MSNs in the DMS (200-300 μW, 10 Hz, pulse width). For NpHR, green light was generated by a 532 nm laser (Shanghai Laser & Optics Century Co. LTD) and to inhibit dmPFC cell bodies and projection fibers (bilaterally) in the DMS (5 mW, constant). A Maser-8 (A.M.P.I.) pulse generator controlled by TTL pulses from Ethovision (Noldus) software was used to drive the laser throughout the trial. Laser output was delivered to the animal via an optical fiber (0.39 NA, 200 μm, Thorlabs) connected to a 1x1 fiber optic rotary joint (Doric Lenses) followed by another optical fiber (0.37 NA, 200 μm, Doric Lenses) which was coupled to the cannula through a ceramic sleeve (Thorlabs).

*dmPFC cell body ChR2 and NpHR, dmPFC to DMS projection ChR2*

Each trial consisted of a 25-minute laser stimulation paradigm. This consisted of a 5-minute baseline laser off period followed by 10 2-minute on/off alternating epochs.

*dmPFC to DMS projection NpHR*

For this bilateral inhibition, we had a 5-minute baseline followed by 5 minutes laser on, 5 minutes laser off, repeating for a total of 25 minutes per trial (5-minute baseline plus 4 on/off 5-minute epochs).

**Chemogenetic Manipulations**

For the DREADDs experiments we injected .5 mg/kg of CNO hydrochloride (diluted in 0.9% saline, Sigma Aldrich). Control animals were injected with saline (0.9%). Injections were done 10 minutes prior to starting the behavior experiment.

**Ex vivo Electrophysiology**

For ex vivo (slice) electrophysiology experiments, we injected adult D1-tmt mice with AAV-CaMKII-ChR2-eYFP (see above) in the mPFC. 4-6 weeks after surgery, animals were terminally anesthetized with ketamine/xylazine, and transcardially perfused with ice-cold, carbogenated glycerol-based artificial cerebrospinal fluid (aCSF) containing (in mM) 250 glycerol, 2.5 KCl, 1.2 $NaH_2PO_4$, 10 HEPES, 21 $NaHCO_3$, 5 D-glucose, 2 $MgCl_2$, 2 $CaCl_2$. The brain was dissected and glued to a chuck, and submerged in ice-cold, carbogenated glycerol-based aCSF. Coronal slices (300 μm) containing the striatum were cut using a vibrating microtome (Leica) and immediately transferred to a chamber containing warmed (34 °C ) carbogenated aCSF containing (in mM) 125 NaCl, 26 $NaHCO_3$, 2.5 KCl, 1.25 $NaH_2PO_4$, 12.5 D-glucose, 1 $MgCl_2$, 2 $CaCl_2$. After incubation for 60 minutes, slices were stored in carbogenated aCSF at room temperature until used for recordings.

For recordings, slices were transferred to a stage-mounted chamber on an Olympus BX51 microscope. Slices were superfused with warmed carbogenated aCSF (31-33 °C) throughout. The DMS was identified at low power, and the area of greatest terminal field ChR2-YFP expression was chosen for subsequent whole-cell recordings. In a given field under high power, medium-sized ovoid cell bodies were targeted using differential interference contrast (DIC) optics. The presence or absence of tdTomato fluorescence was used to determine if an individual cell body belonged to a direct pathway (D1) or indirect pathway (D2) neuron. Since tdTomato-negative neurons could include striatal interneurons, we excluded neurons with physiological features of interneurons (membrane tau decay of <1 msec). D1 and D2 neurons were patched in nearby serial pairs, in randomized order. All whole-cell recordings were acquired (filtered at 5 kHz) using a Multiclamp 700B amplifier (Molecular Devices) and digitized (10 kHz) using an ITC-18 A/D board (HEKA). Igor Pro 6.0 software and custom acquisition routines (mafPC, courtesy of Matthew A. Xu-Friedman) were used to acquire and analyze the data.

Neurons were patched in the whole-cell voltage-clamp configuration using borosilicate glass electrodes (3-5 MΩ). To record EPSCs, we used a cesium methanesulfonate-based, low chloride internal solution containing (in mM) 120 $CsMeSO_3$, 15 CsCl, 8 NaCl, 0.5 EGTA, 10 HEPES, pH = 7.3. The internal chloride concentration was calibrated such that the reversal potential of $GABA_A$-mediated (disynaptic) IPSCs was -70 mV (thus currents recorded at -70 mV were predominantly glutamatergic in origin). Experiments were performed in picrotoxin to pharmacologically isolate EPSCs. mPFC-derived EPSCs were measured at -70 mV holding potential, evoked using brief (3 msec) full-field blue (473 nm) light pulses delivered by a TTL-controlled LED (Olympus) through a ChR2 filter. Light power (473 nm) was set at 1 mW at the objective using a light meter (Thorlabs). EPSC amplitude was defined as the average difference between the baseline holding current (0-100 msec prior to the light pulse) and the peak of the evoked EPSC, averaged over at least five trials (20-second intertrial interval).

**Perfusion/Histology**

Following the conclusion of our behavioral experiments, animals were anesthetized using 5% isoflurane and given a lethal dose (1.0 mL) cocktail of ketamine/xylazine (10 mg/ml ketamine, 1 mg/ml xylazine).

They were then transcardially perfused with 10 mL of 1X PBS followed by 10 mL 4% paraformaldehyde (PFA). They were left in 4% PFA overnight and then transferred to a 30% sucrose solution until slicing. The brains were frozen and sliced on a sliding microtome (Leica Biosystems) and placed in cryoprotectant in a well-plate. Slices were then washed in 1xPBS, mounted on slides (Fisherbrand Superfrost Plus) and air dried (covered). ProLong Gold antifade reagent (Invitrogen, ThermoFisher Scientific) was injected on top of the slices and a cover slip (Slip-rite, ThermoFisher) was placed on top and the slides were left to dry overnight (covered). Viral injection, fiber photometry cannula implant, and optogenetic cannula implant placements were histologically verified on a fluorescence microscope (Leitz DMRB, Leica).

**Statistical Analysis**

Statistical Analysis was done using Prism 7 (Graphpad Software). Unpaired t-test, Paired t-test, Wilcoxon signed-rank, one-way ANOVA with Tukey's correction for multiple comparisons, two-way repeated measures ANOVA with Sidak's correction for multiple comparison was used.

### Supplemental Figure 1

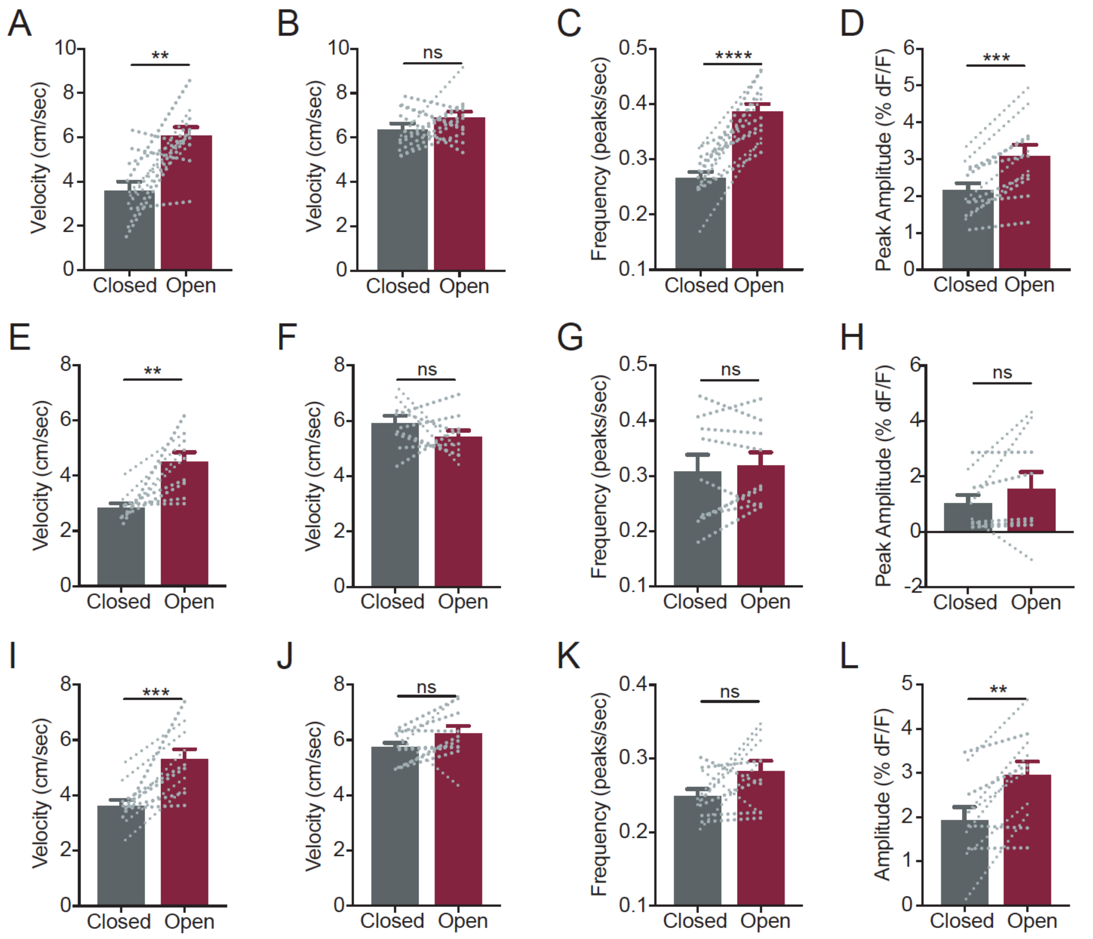


### Supplemental Figure 2

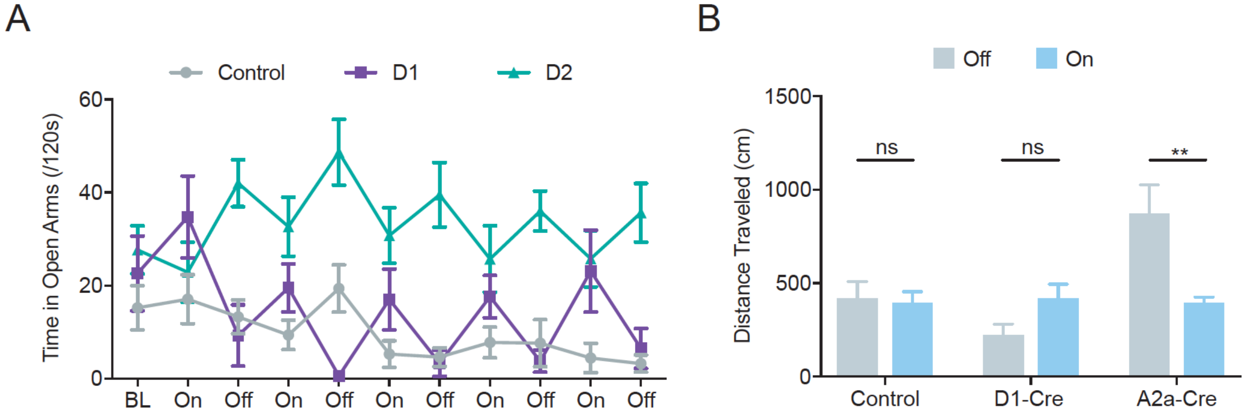

Supplemental Figure 3

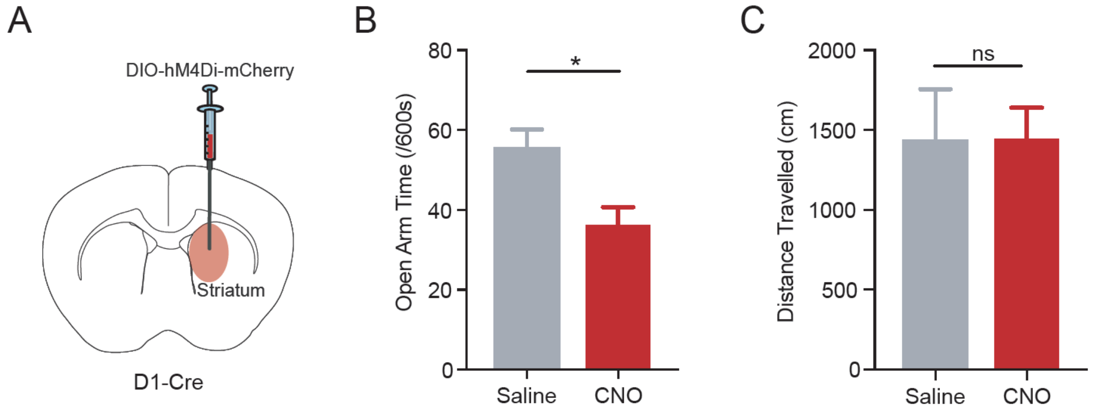

### Supplemental Figure 4

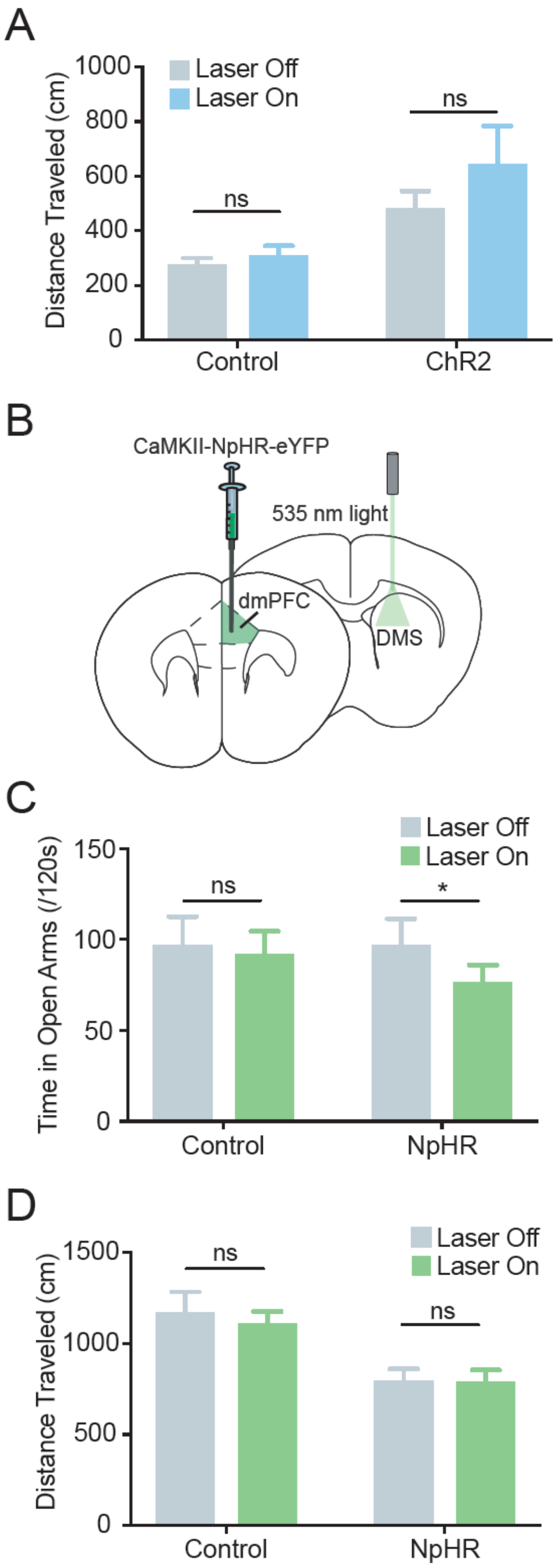

Supplemental Figure 5

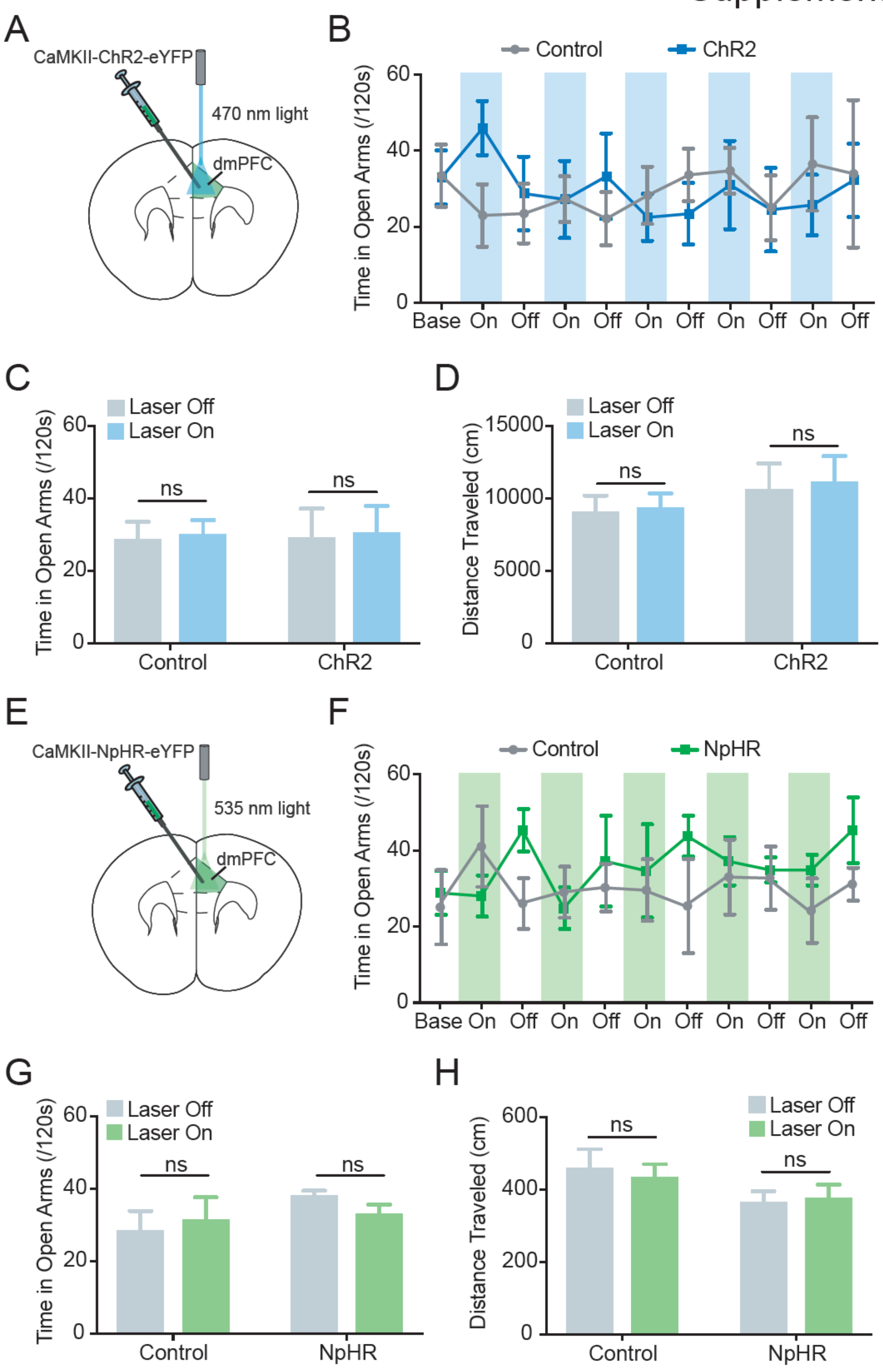

### SUPPLEMENTAL FIGURE LEGENDS

**Figure S1 (related to Figures 1 and 2). Velocity-correcting bulk calcium imaging data does not affect significance of dmPFC and dmPFC projection results.**

**(A)** Whole population dmPFC mice show increased velocity of movement in the open arms than in the closed arms (paired t-test, $p = 0.0032$; $N_{mice} = 11$). **(B)** Once data is velocity corrected (see Methods) there is no longer a difference. (paired t-test, $p = 0.2982$; $N_{mice} = 11$). **(C)** Frequency of calcium transients is significantly higher in the open arms than in the closed arms (velocity-corrected) (paired t-test, $p < 0.0001$; $N_{mice} = 11$). **(D)** Peak amplitude of calcium transients is significantly higher in the open arms than in the closed arms (velocity-corrected) (paired t-test, $p = 0.0001$; $N_{mice} = 11$). **(E-H)** Same as A-D for dmPFC-BLA projection data. **(E)** dmPFC-BLA mice show increased velocity of movement in the open arms than in the closed arms (paired t-test, $p = 0.0038$; $N_{mice} = 9$). **(F)** Once data is velocity corrected (see Methods) there is no longer a difference (paired t-test, $p = 0.2889$; $N_{mice} = 9$). **(G)** No difference in frequency of calcium transients in the open arms than in the closed arms (velocity-corrected) (paired t-test, $p = 0.4533$; $N_{mice} = 9$). **(H)** No difference in peak amplitude of calcium transients in the open arms than in the closed arms (velocity-corrected) (paired t-test, $p = 0.2849$; $N_{mice} = 9$). **(I-L)** Same as A-D for dmPFC-DMS projection data. **(I)** dmPFC-DMS mice show increased velocity of movement in the open arms than in the closed arms (paired t-test, $p = 0.0009$; $N_{mice} = 10$). **(J)** Once data is velocity corrected (see Methods) there is no longer a difference. (paired t-test, $p = 0.1614$; $N_{mice} = 10$). **(K)** No difference in frequency of calcium transients in the open arms than in the closed arms (velocity-corrected) (paired t-test, $p = 0.1060$; $N_{mice} = 10$). **(L)** Peak amplitude of calcium transients is significantly higher in the open arms than in the closed arms (velocity-corrected) (paired t-test, $p = 0.001$; $N_{mice} = 10$).

**Figure S2 (related to Figure 3). Optogenetic stimulation of D1 and D2 MSNs leads to a bidirectional effect on approach-avoidance behavior.**

**(A)** D1-ChR2 mice show a selective increase in time spent in open arms during laser-on epochs across the entire stimulation paradigm (5-minute baseline followed by 2-minute on/off epoch of laser stimulation).

A2a-ChR2 mice show a selective decrease in time spent in open arms during laser-off epochs across the paradigm. Control mice show no modulation of time spent in open arms in response to laser stimulation. **(B)** Control mice and D1-ChR2 mice show no locomotor effect of laser stimulation. A2a-ChR2 show a decrease in distance travelled during laser-on epochs as compared to laser-off epochs (two-way RM ANOVA interaction, $F_{(2,46)} = 5.985$, $p = 0.0049$; Sidak's multiple comparisons, $p = 0.990$ (eYFP, laser on vs. off), $p = 0.5530$ (D1-ChR2, laser on vs. off, $p = 0.016$ (A2a-ChR2, laser on vs. off); $N_{eYFP} = 10$ mice, $N_{D1\text{-}ChR2} = 6$ mice, $N_{A2a\text{-}ChR2} = 10$ mice).

**Figure S3 (related to Figure 3). Chemogenetic inhibition of D1 MSNs causes an increase in avoidance behavior on the EPM.**

**(A)** Schematic of DREADDs experiment showing injection of cre-dependent inhibitory DREADDs (hM4Di) into the DMS of D1-Cre mice allowing for chemogenetic inhibition of D1 MSNs following intraperitoneal cloazapine-n-oxide (CNO) injection (.5mg/kg). **(B)** CNO injected mice show a decrease in time spent in the open arms of the elevated plus maze (EPM) as compared to saline injected mice (unpaired t-test, $p = 0.0421$; $N_{saline} = 3$ mice, $N_{CNO} = 3$ mice). **(C)** CNO injection shows no effect on locomotion (unpaired t-test, $p = 0.9831$; $N_{saline} = 3$ mice, $N_{CNO} = 3$ mice).

**Figure S4 (related to Figure 4). Optogenetic stimulation of dmPFC-DMS neurons has no effect on locomotion while inhibition of dmPFC-DMS neurons increases avoidance behavior in the EZM.**

**(A)** ChR2 mice show no effect of laser stimulation on locomotion. **(B)** Schematic of optogenetic inhibition of dmPFC-DMS neurons. CaMKII-NpHR-eYFP was virally expressed in the dmPFC and a 200 µm optical fiber was implanted in the DMS. Mice were optogenetically inhibited (535 nm light) during exploration of the EZM. **(C)** ChR2 mice show a decrease in time spent in the open arms during laser-on epochs as compared to laser off. Control mice show no modulation of open arm time in laser on versus laser-off epochs (two-way RM ANOVA interaction, $F_{1,19} = 1.911$, $p = 0.1828$; Sidak's multiple comparisons, $p = 0.0221$ (NpHR, laser on vs. off), $p = 0.7989$ (eYFP, laser on vs. off); $N_{NpHR} = 12$ mice, $N_{eYFP} = 9$ mice). **(D)** NpHR and control mice show no effect of laser stimulation on locomotion (two-way RM ANOVA interaction, $F_{1,19} = 0.9647$, $p = 0.3383$; Sidak's multiple comparisons, $p = 0.9824$ (NpHR, laser on vs. off), $p = 0.3020$ (eYFP, laser on vs. off); $N_{NpHR} = 12$ mice, $N_{eYFP} = 9$ mice).

**Figure S5 (related to Figure 4). Optogenetic stimulation and inhibition of the dmPFC as a whole has no effect on approach-avoidance behavior.**

**(A)** Schematic of optogenetic stimulation of dmPFC pyramidal neurons. CaMKII-ChR2-eYFP was virally expressed in the dmPFC and a 200 μm optical fiber was implanted above. Mice were optogenetically stimulated (470 nm light) during exploration of the EZM. **(B)** ChR2 and control mice show no effect of laser stimulation on time spent in open arms during laser-on epochs across the entire stimulation paradigm (5-minute baseline followed by 2-minute on/off epoch of laser stimulation). **(C)** ChR2 and control mice show no modulation of open arm time in laser on versus laser-off epochs (two-way RM ANOVA interaction, $F_{1,9} = 0.0003$, $p = 0.9876$; Sidak's multiple comparisons, $p = 0.8806$ (ChR2, laser on vs. off), $p = 0.8476$ (eYFP, laser on vs. off); $N_{ChR2} = 5$ mice, $N_{eYFP} = 6$ mice). **(D)** ChR2 and control mice show no effect of laser stimulation on locomotion (two-way RM ANOVA interaction, $F_{1,9} = 0.1808$, $p = 0.6806$; Sidak's multiple comparisons, $p = 0.2578$ (ChR2, laser on vs. off), $p = 0.4813$ (eYFP, laser on vs. off); $N_{ChR2} = 5$ mice, $N_{eYFP} = 6$ mice). **(E)** Schematic of optogenetic inhibition of dmPFC pyramidal neurons. CaMKII-NpHR-eYFP was virally expressed in the dmPFC and a 200 μm optical fiber was implanted above. Mice were optogenetically inhibited (535 nm light) during exploration of the EZM. **(F)** NpHR and control mice show no effect of laser stimulation on time spent in open arms during laser-on epochs across the entire stimulation paradigm (5-minute baseline followed by 2-minute on/off epoch of laser stimulation). **(G)** NpHR and control mice show no modulation of open arm time in laser on versus laser-off epochs (two-way RM ANOVA interaction, $F_{1,7} = 2.429$, $p = 0.1631$; Sidak's multiple comparisons, $p = 0.4079$ (NpHR, laser on vs. off), $p = 0.6560$ (eYFP, laser on vs. off); $N_{NpHR} = 4$ mice, $N_{eYFP} = 5$ mice). **(H)** NpHR and control mice show no effect of laser stimulation on locomotion (two-way RM ANOVA interaction, $F_{1,7} = 0.1421$, $p = 0.7174$; Sidak's multiple comparisons, $p = 0.9839$ (NpHR, laser on vs. off), $p = 0.9187$ (eYFP, laser on vs. off); $N_{NpHR} = 4$ mice, $N_{eYFP} = 5$ mice).
